# Compositional Data Analysis is necessary for simulating and analyzing RNA-Seq data

**DOI:** 10.1101/564955

**Authors:** Warren A. McGee, Harold Pimentel, Lior Pachter, Jane Y. Wu

## Abstract

*Seq techniques (e.g. RNA-Seq) generate *compositional* datasets, i.e. the number of fragments sequenced is not proportional to the sample’s total RNA content. Thus, datasets carry only relative information, even though absolute RNA copy numbers are of interest. Current normalization methods assume most features do not change, which can lead to misleading conclusions when there are many changes. Furthermore, there are few real datasets and no simulation protocols currently available that can directly benchmark methods when many changes occur.

We present **absSimSeq**, an R package that simulates compositional data in the form of RNA-Seq reads. We compared **absSimSeq** with several existing tools used for RNA-Seq differential analysis: sleuth, DESeq2, edgeR, limma, sleuth and ALDEx2 (which explicitly takes a compositional approach). We compared the standard normalization of these tools to either “compositional normalization”, which uses log-ratios to anchor the data on a set of negative control features, or RUVSeq, another tool that directly uses negative control features.

Our analysis shows that common normalizations result in reduced performance with current methods when there is a large change in the total RNA per cell. Performance improves when spike-ins are included and used with a compositional approach, even if the spike-ins have substantial variation. In contrast, RUVSeq, which normalizes count data rather than compositional data, has poor performance. Further, we show that previous criticisms of spike-ins did not take into consideration the compositional nature of the data. We demonstrate that **absSimSeq** can generate more representative datasets for testing performance, and that spike-ins should be more frequently used in a compositional manner to minimize misleading conclusions in differential analyses.

**Author Summary:** A critical question in biomedical research is “Is there any change in the RNA transcript abundance when cellular conditions change?” RNA Sequencing (RNA-Seq) is a powerful tool that can help answer this question, but two critical parts of obtaining accurate measurements are (A) understanding the kind of data that RNA-Seq produces, and (B) “normalizing” the data between samples to allow for a fair comparison. Most tools assume that RNA-Seq data is count data, but in reality it is “compositional” data, meaning only percentages/proportions are available, which cannot directly answer the critical question. This leads to distorted results when attempting to simulate or analyze data that has a large global change.

To address this problem, we designed a new simulation protocol called ***absSimSeq*** that can more accurately represent RNA-Seq data when there are large changes. We also proposed a “compositional normalization” method that can utilize “negative control” features that are known to not change between conditions to anchor the data. When there are many features changing, this approach improves performance over commonly used normalization methods across multiple tools. This work highlights the importance of having negative controls features available and of treating RNA-Seq data as compositional.

## Introduction

High-throughput methods, including RNA-Seq, are frequently used to determine what features—genes, transcripts, protein isoforms—change in abundance between different conditions [1]. Importantly, though, researchers ultimately care about the absolute abundance of RNA transcripts. In other words, is there a change in the number of RNA molecules in a cell when the conditions change? However, current techniques are limited to reporting relative abundances of RNA molecules: the proportion of fragments generated by a sequencer that contain a given sequence [2-5]. This means that RNA-Seq is inherently compositional data, where relative proportions are the only information available, yet those are being used to draw conclusions about the absolute abundance of features [2-5] (see Note 1 in **Supporting Information**). Several studies have raised the alarm on ways in which interpretation of the results can be distorted if RNA-Seq data are not properly treated as compositional [2-6].

The first statistical problem in an RNA-Seq analysis lies in determining the origin of the fragments generated. There are two classes of tools available to solve this problem: (1) tools that use traditional alignments to determine the exact genomic location (**tophat2**, **bwa**, **STAR**, **HISAT2**, etc.) (reviewed in [7]); there are other tools that take these traditional alignments and estimate exon-, transcript-, or gene-level expression levels (reviewed in [8]); (2) tools that probabilistically estimate transcript sets that are compatible with producing the corresponding fragments using pseudoalignment and quantify the levels of transcript expression (**kallisto**, **salmon**, **sailfish**) [9-11].

The second statistical problem, the focus of this paper, is to compare the differences in samples collected under different experimental conditions (e.g. comparing cancer cells with control cells; comparing wild-type cells with mutant cells). We will refer to this second step as “differential analysis.” A number of tools are available for differential analyses (**DESeq2**, **edgeR**, **limma-voom**, etc), using continuous or count data (reviewed in [1,12]). One recently developed tool, **sleuth**, utilizes the bootstraps produced by the quasi-mapping tools to estimate the technical variation introduced by the inferential procedure [13].

It is a recognized need to normalize and transform the data before conducting differential analyses. Multiple strategies have been developed to meet this need, including quantile normalization [14], the trimmed mean of M-values (**TMM**) method used by **edgeR** [15], the median ratio method used by **DESeq** and **DESeq2** [16], and the **voom** transformation used by **limma** [17]. In addition, multiple units are used when modeling and reporting RNA-Seq results [18], including counts [16,19], CPM [17], FPKM [20], and TPM [21]. Importantly, all of these strategies, even those that are focused just on the counts for each feature, utilize units that are really proportions, which belies the fact that RNA-Seq data are compositional [5,22] (see Note 1 in **Supporting Information**). Furthermore, all of these normalization strategies assume that the total RNA content does not change substantially across the samples (see [23] for a review). This assumption allows users to leap from the inherently relative information contained in the dataset to the RNA copy number changes in the population under study without quantifying the actual RNA copy numbers. However, there are biological contexts where this assumption is not true [4,24,25], and it is unclear how much change can occur before distorting results when the datasets are not considered as compositional during analyses.

If information is available about negative controls (e.g. spike-ins, validated reference genes), then such information could be used to anchor the data. This has been done in several studies, where the use of spike-ins led to a radically different interpretation of the data compared to the standard pipeline [4,24,26]. In one study, the RUVg approach was designed to use this reference information to normalize RNA-Seq data, as part of the RUVSeq R package [27]. There have been recommendations to include spike-ins as part of the standard protocol [22,24,28]. However, Risso et al. observed significant variation in the percentage of reads mapping to spike-ins, as well as discordant global behavior between spike-ins and genes [27]. While spike-ins are often used in single-cell RNA-seq applications, they are not routinely used in bulk RNA-Seq experiments.

John Aitchison developed an approach to compositional data with the insight that ratios (or log-transformed ratios, called “log-ratios”) capture the relative information contained in compositional data [29]. There are three requirements for any analytical approach to compositional data: scale invariance, subcompositional coherence, and permutation invariance (see Note 2 in **Supporting Information**) [30]. It was recently demonstrated that correlation, a widely used measure of association in RNA-Seq analysis, is subcompositionally incoherent and may lead to meaningless results, and an alternative called “proportionality” was proposed [4]. A tool was previously developed to apply compositional data analysis to differential analysis, called ALDEx2 [3]. However, ALDEx2 is not well-suited for utilizing the bootstraps generated by the pseudoalignment tools and is unable to detect any differentially expressed features when there are less than five replicates [31]. Therefore, it is necessary to develop a compositional approach for other tools.

Two tools are commonly used to simulate RNA-Seq: polyester and RSEM-sim [21,32]. These tools require the input of estimated counts per transcript and the expected fold changes between groups. However, without considering the data as compositional, protocols used to simulate RNA-Seq data result in the total read counts being confounded with the condition, such that one condition will have on average a greater depth compared to the other condition (for example, see Supplementary Table S2 of [13]). A protocol that simulates many changing features and at the same time yields similar sequencing depth per sample, is lacking, but could be done using principles from compositional data analysis. This challenge, along with the one above, both motivated the present work.

Here, we present **absSimSeq**, a protocol to simulate RNA-Seq data using concepts from compositional data analysis. This protocol allows us to directly model large global shifts in RNA content while still maintaining equivalent sequencing depths per sample. Further, we developed a normalization approach that uses negative control features (e.g. spike-ins) with log-ratios, which we call “compositional normalization”. We created an extension of sleuth, called **sleuth-ALR**, to use compositional normalization, both to predict candidate reference genes and to normalize the data. We also adapted already available methods to implement compositional normalization for other differential analysis tools. We then used absSimSeq to benchmark performance of differential analysis tools in the setting of either a small or large change to the total RNA content. Within each setting, we compared the current normalization approaches versus either compositional normalization or the RUVg approach with spike-ins.

When there was only a small change in total RNA content, all tested tools had similar performance on simulated data, whereas sleuth, sleuth-ALR and limma had the best performance on real data. However, when there was a large change in the RNA content, either up or down, all tools had much better performance if compositional normalization with spike-ins was used. When analyzing a well-characterized real dataset which had a large change in total RNA content per cell [4,26], only compositional normalization with a validated reference gene was able to capture the overall decrease in RNA transcription. Surprisingly, RUVg had poor performance, and the choice of normalization approach had a greater impact on performance than the choice of tool. Furthermore, both of the concerns about spike-ins raised by Risso and colleagues [27] are actually the expected consequences of how compositional data behaves between samples, though they do raise concerns about the proper protocol for including spike-ins.

In summary, we provide **absSimSeq** as a resource to generate simulated RNA-Seq datasets that more accurately reflect the behavior of real datasets. This will help future development of RNA-Seq analysis when testing performance. Furthermore, our work suggests that using compositional normalization with spike-ins or validated reference genes is essential for differential analyses of RNA-seq data. When such information is missing, it raises major concerns about the limitations of drawing conclusions from the inherently compositional data of RNA-Seq and other “omics” techniques.

## Results

### Simulation of RNA transcript copy numbers, normalization, and performance of different tools

To test how changes in the total RNA content can affect performance of differential analysis tools, we developed **absSimSeq** to simulate RNA-Seq data (**Fig 1**). Because the experimental step of generating a library from samples in a real RNA-Seq experiment generates a compositional dataset, this protocol directly simulates that step, producing simulated compositional data. Using this protocol, we carried out three simulation studies, each having five experiments. In each study, current normalization methods were compared to compositional normalization and **RUVg**. A set of highly expressed spike-ins was used as the set of negative control features for compositional normalization and RUVg (see methods for details).

**Fig 1.**
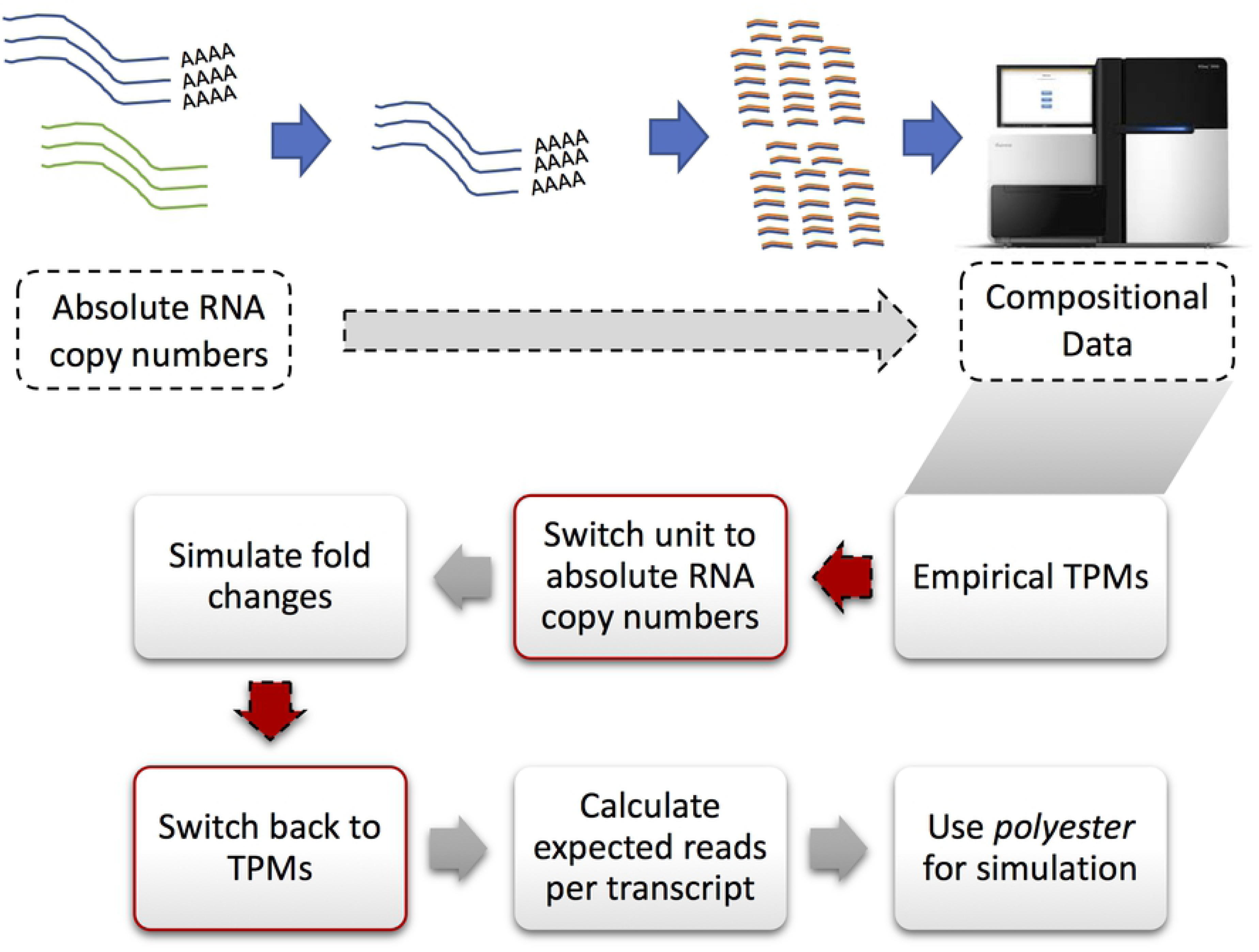
AbsSimSeq, A novel simulation protocol to model compositional RNA-Seq data. All RNA-Seq experiments convert copy numbers per cell to relative abundances because of the selection step and because the depth of sequencing is arbitrary with respect to the total RNA present (top panel; see Note 1 in **Supporting Information**). The **absSimSeq** protocol simulates that conversion process (bottom diagram), with the key, novel steps highlighted in red. It takes the mean empirical relative abundances (in TPM units) from any dataset. It then makes a conceptual leap by assuming those values are copy numbers per cell. Then, the fold changes are simulated on these copy numbers for the experimental condition. After that, in the crucial step of the protocol, the copy numbers are re-normalized back to relative abundances to simulate what happens in the RNA-Seq experiment. From there, the expected reads per transcripts are calculated using relative abundances and the median of the estimated effective lengths from the original dataset. These are then submitted to the polyester R package for a negative binomial simulation.

In the first study (“small”), only a small fraction of features were changed (5% of all transcripts), and the total RNA content was similar between experimental groups (<2% change) (**S1 Table** and **S2 Table**). This study was intended to simulate an experiment that fulfills the assumption of the current normalization methods. Under these conditions, all tools tested performed similarly whether using their current normalization approaches or using compositional normalization (**Fig 2A**; **S2 Fig**, panel A).

**Fig 2.**
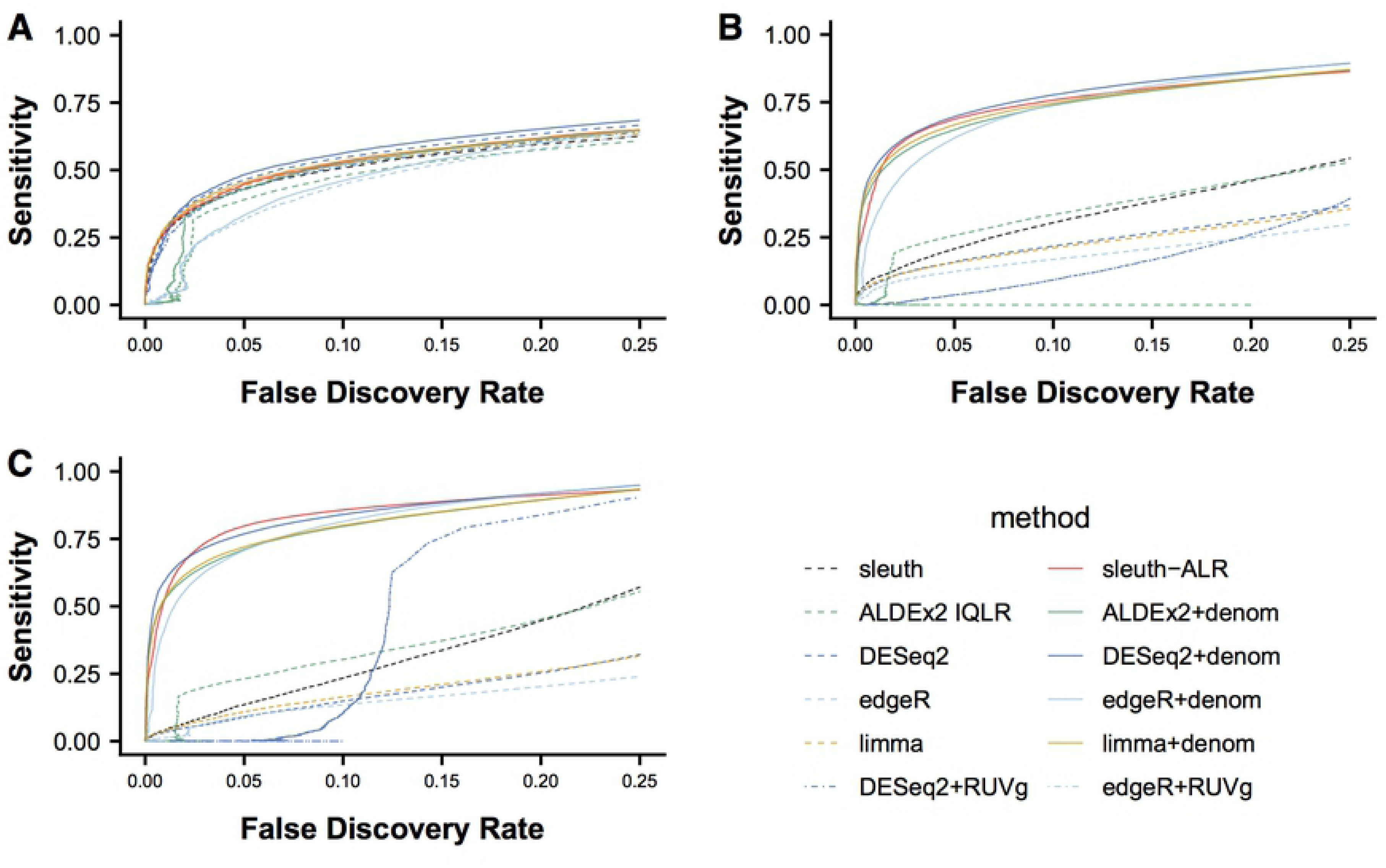
Compositional normalization markedly improves performance when there is a large compositional change. The copy numbers were modeled using the estimated average abundances from the GEUVADIS Finnish women samples (N = 58). Each of the three studies consists of five simulations under specified global conditions: (A) the “small” group has 5% of the transcriptome (∼10K transcripts) simulated as differentially expressed, with the total average copy numbers per cell in each group roughly equal, (B) the “down” group has 20% of the transcriptome (∼40K transcripts) simulated as differentially expressed, with 90% of the transcripts down-regulated, and (C) the “up” group has the same number of transcripts changing as the “down” group, but 90% of the transcripts are up-regulated. The compositional normalization methods (solid lines) used a set of highly expressed spike-ins to illustrate, especially in the “down” and “up” groups, what happens to the performance of tools when there is a large shift in the global copy numbers per cell. Average false discovery rate across the simulations within each group (n = 5) is shown on the x-axis, and average sensitivity is shown on the y-axis. The FDR range between 0 and 0.25 is shown. Note that edgeR+RUVg is not shown because it always had an FDR above 0.25. See **S2 Fig** for the full range.

In the second set of studies (“down” and “up”), many features were differentially expressed (20% of all transcripts), resulting in a large change in the composition, with the total RNA content decreased by ∼33% or increased ∼2.8-fold, respectively (**S1 Table** and **S2 Table**). Under such conditions, compositional normalization led to greatly improved performance for all tools compared to these tools using their current normalization methods (**Fig 2B-C**; **S2 Fig**, panels B-C). In contrast to current normalization methods and compositional normalization, the **RUVg** approach from **RUVSeq** resulted in the worst performance for edgeR and DESeq2 in all three studies (**Fig 2**), even though it used the same set of spike-ins as compositional normalization.

It is worth noting that, among the tools tested, sleuth and ALDEx2 performed the best when there were large compositional changes in the data (**Fig 2B-C**); ALDEx2 uses the IQLR transformation, which is a compositional approach designed to be robust in the presence of large changes to the composition. We also observed that for each transformation used by ALDEx2, it performed almost identically regardless of statistical test used (**S3 Fig**). Finally, sleuth-ALR had similar performance whether TPMs or estimated counts were modeled, or if the Wald test or the likelihood ratio test was used (**S4 Fig**).

### Performance is not degraded by significant variation in individual spike-ins

A previous study reported that spike-ins had significant variation [27], raising a concern about their utility for normalization. In particular, they observed significant variation both between and within groups. In our simulated data, spike-ins were modeled similarly as other features, with over-dispersed variation, drawing from a negative binomial distribution. When estimating the percentage of transcript fragments mapping to spike-ins per sample, we also detected significant variation across samples (**Fig 3**). Importantly, in the “up” and “down” studies, there were systematic differences between groups, similar to what was observed in the MAQC-III study and in the zebrafish study previously analyzed [27]. These systematic differences between groups are expected given the compositional nature of the data (see Note 3 in **Supporting Information** and Discussion). We further observed a large spectrum of estimated fold changes across individual spike-ins, with a systematic asymmetry in the distribution of fold changes in the experiments from the “up” and “down” studies (**S5 Fig**). Despite the significant variation of spike-ins, individually and collectively, spike-ins led to greatly improved performance when used for normalization (**Fig 2**). Consistent with previous work [33,34], our results suggest that the ratio information contained in spike-ins are collectively robust to variation, and that spike-ins can be used for sample-wise normalization.

**Fig 3.**
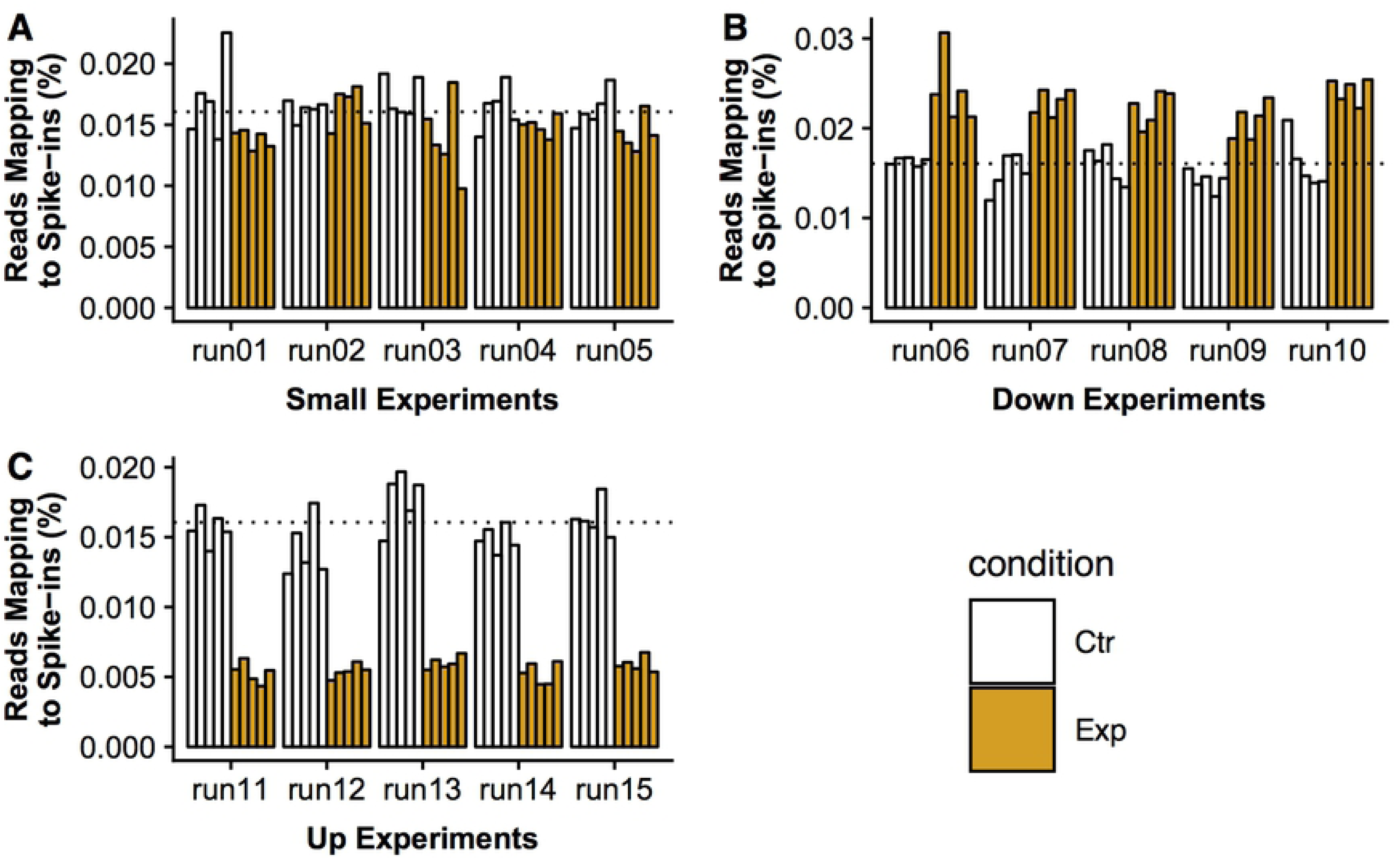
Spike-ins have significant within-group and between-group variation, despite improved performance when used for normalization. Previous work expressed a concern about variation observed in spike-ins between samples. In each experiment, the 92 ERCC spike-ins from Mix 1 were simulated to have no change in copy numbers between the two conditions, as well as to have over-dispersed variation between samples, drawing from a negative binomial distribution. Plotted here is the percentage of all fragments that map to spike-ins, compared to the total number of fragments from the sample, in **(A)** the “small” study, with <2% change in the total RNA in each condition; **(B)** the “down” study, with a ∼33% decrease in total RNA in the experimental condition; and **(C)** the “up” study, with a ∼2.8-fold increase in total RNA in the experimental condition. The dotted line represents the expected percentage of fragments mapping to spike-ins in the control group. Across all experiments, there is significant within-group variation; in the “down” and “up” studies, there is also significant between-group variation. The latter is to be expected given the compositional nature of the data (see Note 3 in **Supporting Information**).

### Sleuth-ALR has best self-consistency and negative control performance among compositional normalization methods

To confirm that compositional normalization performs similarly to current methods in the context of real data, we repeated the analyses using data from the original study on sleuth [13]. The first test was the “self-consistency” test using data from [35]. We reasoned that a tool should provide consistent results from an experiment, whether a few samples per group are sequenced (in this case, n = 3 per group), or more samples per group are used (n = 7-8), as measured by the “true positive rate” (TPR) and “false discovery rate” (FDR). In this experiment, true positives were defined as hits identified in both the smaller and larger datasets, and false positives were defined as hits identified by the smaller dataset but not by the larger dataset. Among all tools using compositional normalization, sleuth and sleuth-ALR with Wald test showed the best balance between the TPR and FDR (**Fig 4**; **S6 Fig**). Limma-voom and sleuth/sleuth-ALR with the likelihood ratio test had the lowest FDR at the cost of a lower TPR. DESeq2 and edgeR both had higher FDR, and on average slightly lower TPR compared to sleuth-ALR. In contrast, the Welch and Wilcoxon statistics in ALDEx2 were unable to identify any hits in any of the “training” datasets, consistent with a recent benchmarking study [31] (data not shown), suggesting that they have greatly reduced power with less data (N = 3 samples per group). ALDEx2’s “overlap” statistic was able to identify hits, but this led to the worst consistency (i.e. highest false discovery rate) among the tools tested. Finally, while DESeq2 with RUVg had similar performance to DESeq2 with compositional normalization, edgeR with RUVg had among the worst consistency.

**Fig 4.**
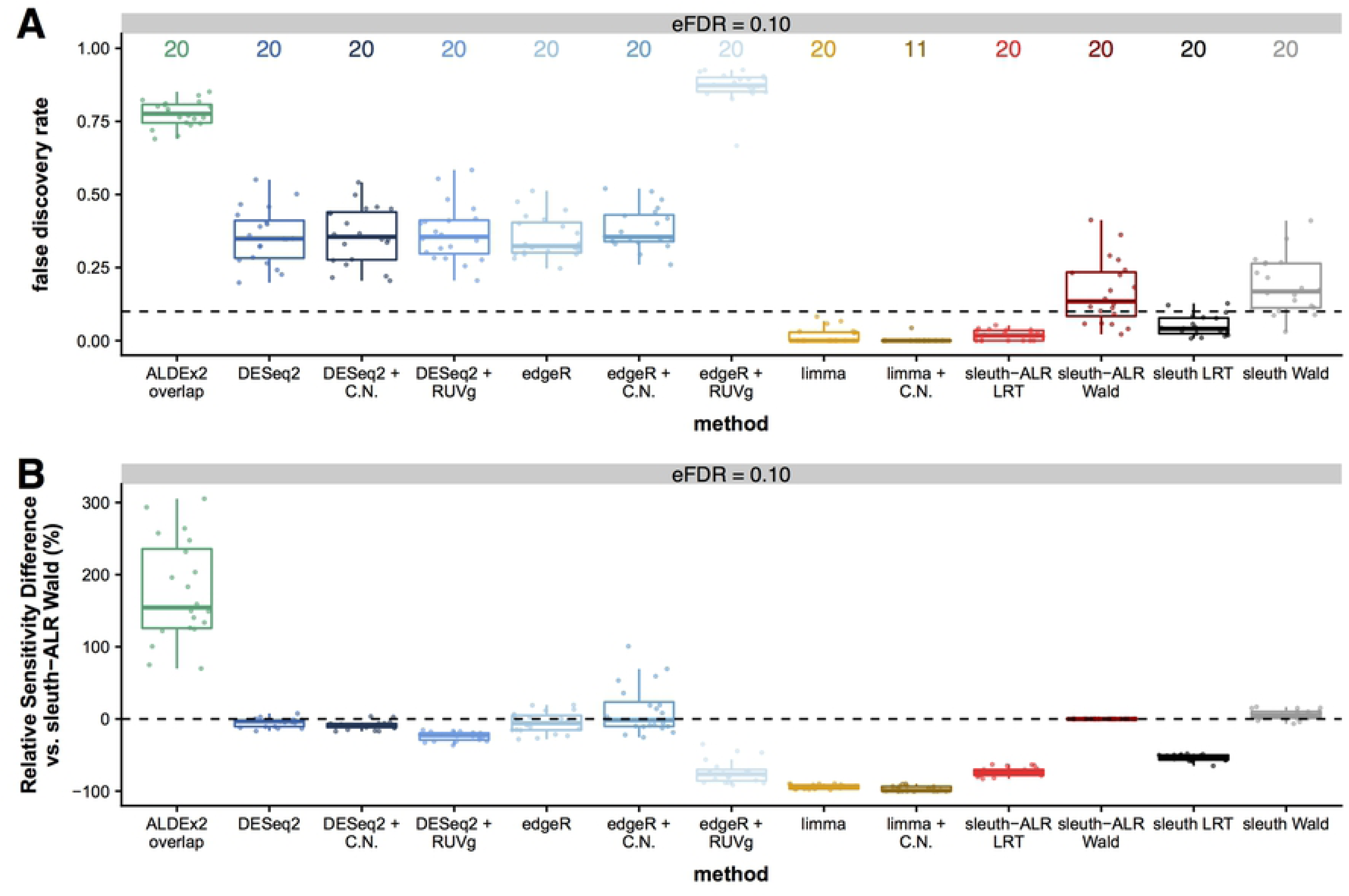
sleuth-ALR Wald has best balance of self-consistency between less and more data from same dataset. Depicted is the Bottomly et al self-consistency test at the isoform level, with **(A)** the false discovery rate at three specified levels, and **(B)** the relative sensitivity as compared to sleuth-ALR with the Wald test. This extends the test from the original sleuth paper [13]. A large dataset is split into a small “training” dataset (3 samples per group), and larger “validation” datasets. A “false discovery” in this test is defined as a hit identified in the “training” dataset but not in the larger “validation” dataset at the given FDR level, and a “true positive” in this test is a hit identified in both datasets at that FDR level. A tool performs well in this test if it can identify the same hits with less data, as well as control the “false discovery rate” at the specified FDR level. The full dataset was split twenty times. Note that the number above each tool in panel A is the number of “training” datasets out of twenty that identified at least one hit at the specified FDR level. See **S6 Fig** for the results at the FDR levels of 0.01 and 0.05.

Next, we tested the performance of compositional normalization on a negative control dataset, where there are no expected differentially expressed features. We repeated the null resampling experiment from the sleuth paper [13] using the GEUVADIS Finnish women dataset (n = 58) [36]. Six samples were randomly selected (stratified by lab to minimize technical variation) and split into two groups, with the expectation of finding no hits. We found that sleuth-ALR with the likelihood ratio test performed similarly to sleuth and limma-voom (median number of false positives < 5) (**Fig 5**), and sleuth-ALR with the Wald test also showed good false positive control (median number of false positives = 10). In contrast, DESeq2 and edgeR with compositional normalization showed higher numbers of false positives (median of 71 and 66, respectively), and the “overlap” statistic for ALDEx2 showed the highest number of false positives (median of >5000 at the 0.1 FDR level).

**Fig 5.**
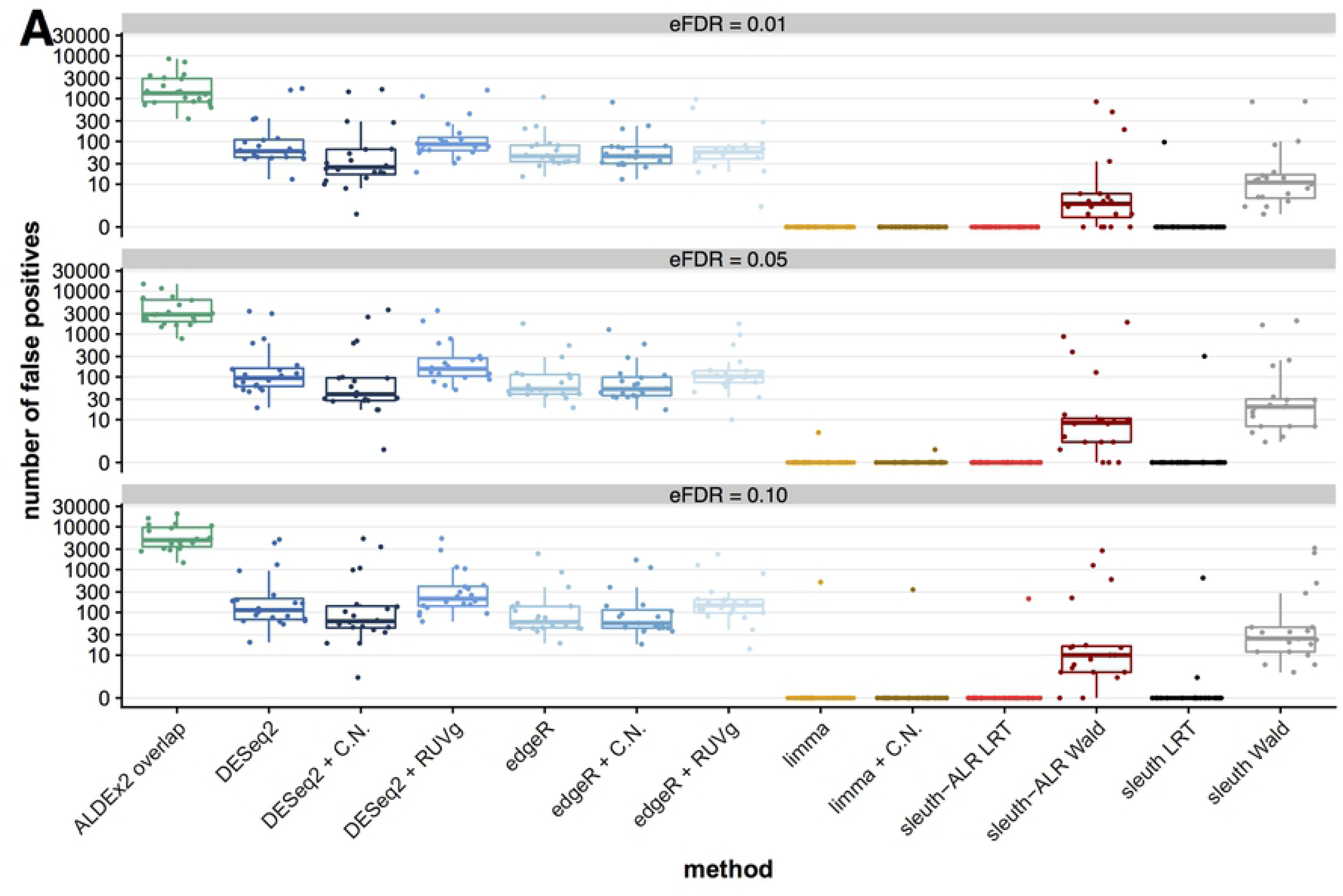
sleuth-ALR and limma perform best on the GEUVADIS null dataset. Depicted is the null experiment at the isoform level **(A)**. This also extends the test from the original sleuth paper [13]. The data were from the lymphoblastoid cells of 58 Finnish women, a relatively homogeneous population, taken from the GEUVADIS project [36]. Data from six women were resampled from the larger dataset, stratifying by lab to minimize technical variation, and then randomly split into two groups to simulate a “null experiment”. The number of false positives, defined as any hits, are reported here based on twenty rounds of resamplings. A tool performs well in this experiment by minimizing the number of hits reported. ALDEx2 used the IQLR transformation; all “C.N.” methods and sleuth-ALR used compositional normalization; all “RUVg” methods used RUVg for normalization.

### Performance of compositional normalization on a dataset with a global decrease in transcription

To compare different tools in a real dataset with a large compositional change, we used the “yeast starvation dataset” [26]. In this dataset, yeast cells were starved of a nitrogen source, inducing them to enter a reversible quiescent state without active cell division [37]. Absolute copy numbers per cell were estimated for each mRNA by being normalized to a collection of 49 mRNAs that were quantified using NanoString nCounter [38]. Our re-analysis clearly shows a large global decrease in RNA content (**Fig 6A**), with ∼95% of genes decreasing in copy numbers per cell in the starvation group versus control, confirming that the dataset has a large compositional change (**Fig 6B**). On the contrary, analyses using previously developed methods failed to identify this pattern of changes in gene expression, only reporting equivalent numbers of hits up- and down-regulated transcripts (**Table 1**).

**Table 1.** Only compositional normalization (C.N.) accurately reflects global decrease in the yeast starvation study. This table shows the number of hits identified by each tool using default settings and kallisto-calculated estimated counts and abundances. “Sleuth-ALR trend” used rqc1 (Pombase: SPAC1142.01) as a denominator; this gene had the most consistent abundance (TPM value) across all samples. The compositional normalization methods (all tools in red: “ALDEx2 ALR overlap”; “sleuth-ALR”; all “+C.N.” tools) used opt3 (Pombase: SPCC1840.12) as a denominator; this gene was considered a “validated reference gene”. “RUVg” for edgeR and DESeq2 (in blue) also used opt3 as a negative control gene. Only compositional normalization methods, using opt3, were able to accurately reflect the severe global decrease observed in the data, as shown by the number of genes showing down-regulation of the absolute counts. Note that ALDEx2 Welch and Wilcoxon statistics yielded <5 significant hits.

| Tool | Up-Regulated Genes | Down-Regulated Genes |
| --- | --- | --- |
| **ALDEx2 ALR overlap** | **719** | **4496** |
| ALDEx2 IQLR overlap | 2716 | 2677 |
| DESeq2 | 2424 | 2415 |
| **DESeq2 + C.N.** | **632** | **4344** |
| **DESeq2 + RUVg** | **698** | **1001** |
| edgeR | 2621 | 2536 |
| **edgeR + C.N.** | **582** | **4290** |
| **edgeR + RUVg** | **109** | **90** |
| limma | 2603 | 2527 |
| **limma + C.N.** | **573** | **4332** |
| sleuth | 2529 | 2554 |
| sleuth-ALR (trend) | 2692 | 2225 |
| **sleuth-ALR** | **614** | **4356** |
| **Absolute Counts** | **522** | **5751** |

**Fig 6.**
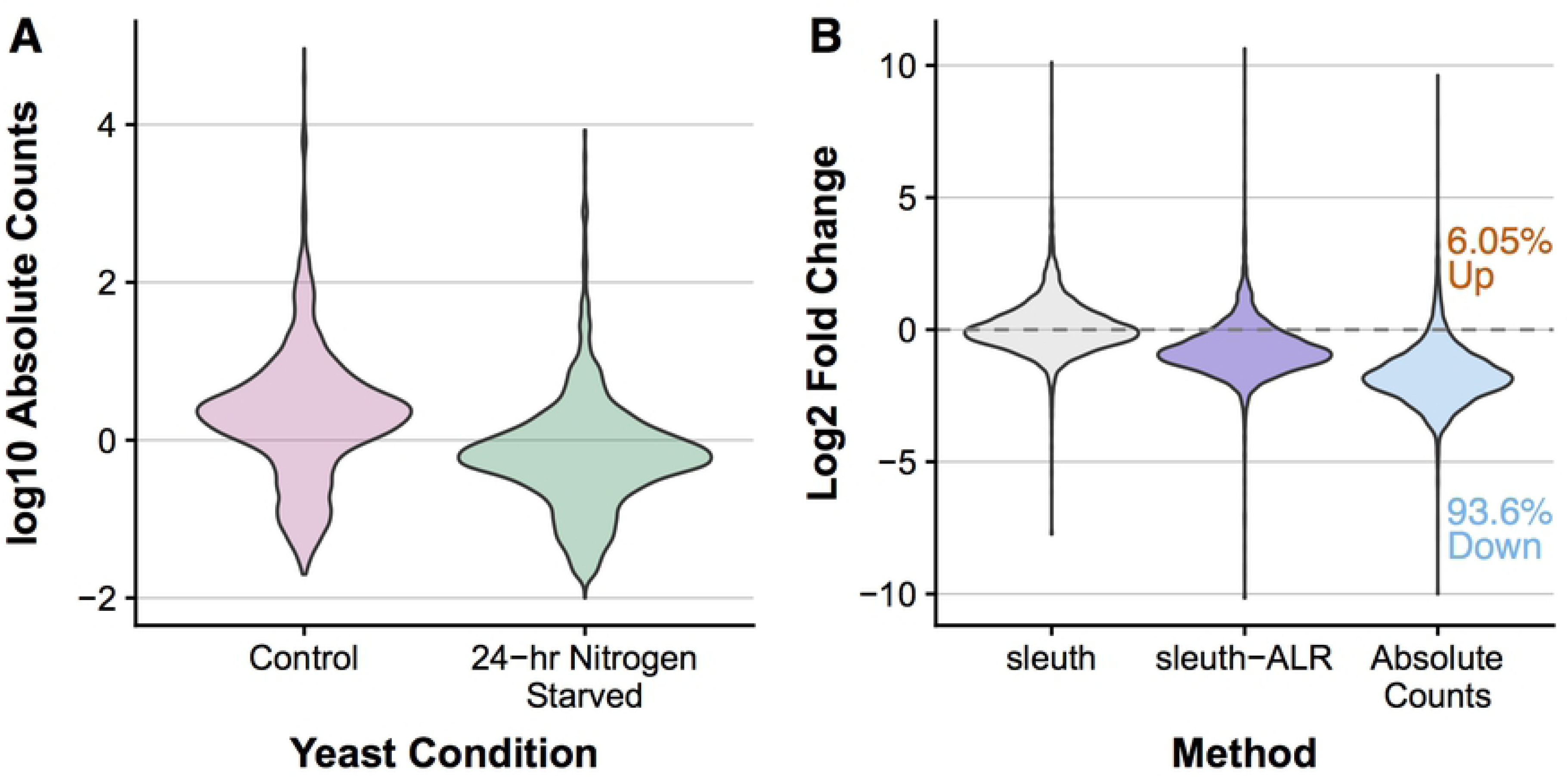
A yeast starvation study shows a large global decrease in RNAs. **(A)** A violin plot showing the distribution of absolute counts in control yeast cells (pink) and yeast cells starved of their nitrogen source for 24 hours (green). The data were from Marguerat et al [26]. The absolute counts were estimated by normalizing RNA-Seq data to a panel of reference genes whose copy numbers were quantified using the NanoString nCounter assay. As can be observed, there is a global decrease in the RNA present. **(B)** a comparison of log2 fold changes calculated using a standard RNA-Seq pipeline (the example shown here in gray is kallisto + sleuth), and the log2 fold changes calculated using compositional normalization (sleuth-ALR, shown in purple) or directly from the estimated absolute counts (in blue). As shown, the vast majority of genes were observed to be downregulated when estimating from the absolute counts. The standard RNA-Seq approach misses this global shift, but compositional normalization is able to identify it.

Next, we examined the importance of using negative control features with compositional normalization. We first analyzed the data using the gene with the most constant proportion across all samples (as measured by coefficient of variation), rqc1 (Pombase: SPAC1142.01). In this context, compositional normalization missed the global pattern of down-regulation; instead, it reported a similar numbers of hits compared to other tools using current normalization methods, both up-regulated and down-regulated (**Table 1**). We then selected a gene with approximately constant expression (opt3, Pombase: SPCC1840.12) as the denominator for compositional normalization. Compared to current methods, compositional normalization using a validated reference gene was able to identify the global decrease in transcription observed by previous analyses of the data [4,26] (**Table 1**). All tools tested (ALDEx2, DESeq2, edgeR, limma-voom, and sleuth-ALR) were able to identify a similar number of hits when using compositional normalization. In contrast, while RUVg was also given the same validated reference gene, surprisingly it had greatly reduced power and was unable to capture the global pattern of down-regulation.

## Discussion

Compositional normalization performed similarly to current normalization methods when there was only a small change to total RNA (**Fig 2A; Fig 4; Fig 5**). This similarity is expected because, in this scenario, it is valid to assume that most features are not changing. However, when there was a large change in global RNA, compositional normalization using negative control features (spike-ins) had much better performance compared to the current normalization methods, for both simulated data (**Fig 2B-2C**) and real data (**Table 1**). In this case, the assumption held by the current methods is violated, and this is likely what greatly reduced their performance. Further, although the IQLR transformation in ALDEx2 was designed to be robust to large changes in global RNA, it only modestly improved performance. This indicates that at least some of the features it selected and assumed to be unchanging were indeed changing, in both the simulated data and the real data.

The worse performance of current normalization methods is likely related to how fold changes behave in the absolute case versus the relative case. Current normalization methods assume that most features are not changing; one could equivalently assume that the total RNA per cell is unchanged [23]. Thus, if the total RNA content changes, current methods will anchor the data on whatever this global change is. This results in a shift in the observed distribution of fold changes (see Supplementary Figure S16 of [4]). Extreme changes will still be observed to have the same direction, but there will be a group of features that are changing less dramatically than the global change that will be observed to have the wrong sign, and many unchanged features will appear to be changing.

Interestingly, the choice of normalization method had a much greater impact on performance than the choice of differential analysis tool, validating a finding from an older study [14]. This was true both for the simulated data (**Fig 2B-2C**) and for the yeast dataset (**Table 1**). In both cases, compositional normalization clearly outperformed current methods when analyzing a dataset with substantial changes to the total RNA content, whether simulated or real. However, this was only true when negative control features were used (**Table 1**). This indicates that the choice of tool is much less important than the choice of normalization and the availability of negative control features (spike-ins, validated reference gene) to properly anchor the data.

Surprisingly, RUVg had poor performance in both the simulated data and the experimental yeast dataset. It is unclear why this occurred, other than the likely possibility that it treats the data as count data rather than compositional data. More work would need to be done to see if RUVg could be modified to more accurately capture the global trend in the data from negative control features.

In our simulations, the total RNA content was either decreased by 33% or tripled, respectively. The overall change in the “up” study was less extreme than what was observed after c-Myc overexpression [24,39]. In that context, the researchers found a general transcriptional activation that was not captured by the traditional analysis of the RNA-Seq data, and required cell number normalization using spike-ins to see the overall trend of increasing gene expression; the total RNA content increase observed by RNA-Seq was ∼5.5-fold (see Table S2 of [24]). The overall change in the “down” group was less extreme than that observed after the Marguerat et al dataset, which observed an 88% decrease total RNA content when using normalized RNA-Seq data. How often large shifts occur in real datasets is unclear because of how infrequently spike-ins or validated reference genes are used when generating data. Future work should determine more carefully how drastic composition changes need to be before performance starts to degrade for methods which assume that most features are not changing.

### Results from Bottomly et al. self-consistency test and GEUVADIS null experiment

Sleuth-ALR had the best self-consistency (**Fig 5**), and sleuth-ALR and limma had the best performance in the negative control dataset (**Fig 6**). ALDEx2 was unable to identify any hits using three samples per group with the standard statistical methods (Wilcoxon and Welch), and its “overlap” statistic showed a very high FDR, indicating that its results were inconsistent between the “training” and larger “validation” datasets. This indicates that ALDEx2 may not perform well when there are few replicates per group. While this manuscript was in preparation, a recent benchmarking study came to the same conclusion [31]. This behavior is likely due to the fact that the algorithm of ALDEx2 does not include any shrinkage of the variance. Variance shrinkage has been demonstrated to improve performance when there are few replicates [17,40,41]. Interestingly, though, all three statistics used by ALDEx2 had similar performance on the simulated data (**S3 Fig**), and the “overlap” statistic identified a similar set of hits in the real dataset as other compositional normalization methods (**Table 1**), suggesting that the “overlap” statistic may have utility in small datasets despite poor self-consistency or poor control of false positives in a negative control dataset. Future work could explore how to improve ALDEx2 performance for smaller datasets.

### The lack of real datasets with verified global changes

It was difficult to identify an example of a real dataset, with clear-cut evidence of substantial changes to the total RNA content, that was also amenable to re-analysis using our pipeline. We were unable to re-analyze the previous data measuring the impact of c-Myc overexpression [24] because the RNA-Seq dataset did not have technical or biological replicates. We were also unable to re-analyze the selective growth assay used in the ALDEx2 paper [3] because the raw data, which is necessary for our pipeline, was not publicly available. Other datasets have used spike-ins, but had no other confirmatory data on the absolute copy numbers to confirm if the spike-ins accurately captured the global trend or not. This dearth of bulk RNA-Seq datasets with verified global changes speaks to how much the problem of neglecting to treat bulk RNA-Seq as compositional data has gone unrecognized in the community.

### How to choose a denominator for compositional normalization and interpret the results

When using compositional normalization, regardless of which denominator is chosen, the interpretation of differential expression and fold-changes is “the change of feature X with respect to the denominator”. Although all transformations are permutation invariant and therefore any chosen denominator will produce mathematically equivalent results [29], the choice of denominator has important implications for the interpretation of the results and for the downstream validation experiments.

If an experimenter has information about absolute copy numbers per cell in their experiment, they can readily use that information with compositional normalization. For example, if spike-ins are included proportional to the number of cells, as recommended in the c-Myc study [24], those spike-ins can be used as the denominator. If one or more reference genes are validated, as was done with the Yeast Starvation study [26], then a reference gene known to be approximately constant under the experimental conditions can be used. In principle, if qPCR is used to validate differential analysis results in this scenario, a predicted reference gene after using spike-ins or the validated reference gene used for compositional normalization would be the best choice for a reference gene.

What about experiments that do not have spike-ins or validated reference genes? Spike-ins have only slowly been adopted as a part of *Seq protocols [28]. It has further been extensively documented that reference genes are frequently not properly validated [42], and that expression of commonly used reference genes could change dramatically under certain circumstances [43,44]. There have been several techniques to identify reference genes using RNA-Seq data [45-47]. Importantly, these techniques all find a feature that has an approximately constant proportion throughout all of the samples. However, researchers are usually attempting to identify a reference gene with approximately constant absolute copy numbers per cell throughout. In order to draw this conclusion, the techniques must make the same assumption that standard RNA-Seq analysis tools make, i.e. that the global RNA content remains constant in all samples, or that only a few features are differentially expressed. If many features are changes, features identified by these tools will only reflect the global change (up or down), rather than being approximately constant in absolute copy numbers per cell.

None of the compositional normalization methods solve this problem (for an example, see sleuth-ALR with the “trend” feature compared to the other methods in **Table 1**), because no tool can *in principle* solve this problem without access to external information. As described in a recent review article [5], no approach can formally recapitulate the absolute data, and only approaches that are using truly constant features can adequately anchor the data to accurately estimate the true changes in the absolute data. In most datasets without spike-ins or validated reference genes, it is unknown if there is a significant change in the total RNA per cell. Thus, all that is left is how one feature behaves relative to another feature.

When one feature is used, there is a clear advantage to compositional normalization versus current methods because there is a clear interpretation of the results (i.e. how features are behaving relative to this feature), and because there would be a clear choice of reference gene for any qPCR validation downstream (for example, spp1 would be used in the yeast starvation study). Any other choice for qPCR reference gene would likely yield discordant results. Importantly, though, identifying a feature with approximately constant proportion, in the absence of information about the overall changes in RNA content, can still help experimenters identify important biology. This is analogous to the approach taken by Gene Set Enrichment Analysis (GSEA) [48]. Its “competitive” null hypothesis leads to the identification of gene sets or pathways that are behaving differently with respect to the general trend of expression changes across the whole genome [49]. GSEA’s approach has led to uncovering interesting biology in the past, as demonstrated by how frequently it has been used and cited.

In the context of RNA-Seq differential analysis, most datasets will be restricted to this option, and thus experiments will be forced to sacrifice knowledge about the absolute copy numbers for an interpretation of the data anchored to whatever the global change is. This should alarm researchers conducting these experiments to recognize the limitations of the current methodologies. This should also push the community to call for technical innovation and standardization that will make more widespread both the use of spike-ins for normalization, and the validation of reference genes specific to the experiment at hand. Furthermore, this issue regarding the compositional nature of the data is not limited to RNA-Seq, but to many if not all high-throughput (“omics”) techniques (see Note 4 in **Supporting Information**).

### Concerns about the utility of spike-ins

The authors of RUVg [27] made two observations that raised concerns about the utility of spike-ins. However, when interpreting the spike-in abundances through the lens of compositional data analysis, the observed behavior of spike-ins is precisely what would be expected (See Note 3 in **Supporting Information**). In particular, systematic variation between conditions in the spike-in abundances is expected if there is a global change (**Fig 3**; **S5 Fig**; **S4 Table**). This was observed in the yeast dataset, with the validated reference gene have a greatly increased abundance in the nitrogen starved cells versus control cells.

However, their other observation raises valid concerns about the current protocol for using spike-ins. They observed a global discrepancy between spike-ins and the rest of the genes when comparing two control libraries to each other (see Figure 4d of [27]). This could be partially explained by dropout effects, but is most likely due to differences in non-poly-adenylated RNA expression (especially rRNA) between the samples. The way that the spike-ins were added in their experiment (adding an equal amount to approximately equal aliquots of the total RNA) causes the spike-ins to also be subject to compositional changes (**S5 Table**). For bulk RNA-Seq experiments, the standard protocol adds spike-ins to equal amounts of RNA after isolation and selection (poly-A selection or rRNA-depletion), but if there are changes in the excluded RNAs, this protocol impedes the ability of the spike-ins to accurately capture the true fold changes of the RNAs under consideration. In contrast, the approach advocated by Lovén et al [24] was to add spike-ins before RNA isolation, in proportion to the number of cells. With this approach, the spike-ins can, in principle, accurately capture the behavior of the genes, even when there are non-poly-adenylated RNA changes (**S6 Table**). There are challenges with using spike-ins in complex tissues [23], and there may be technical biases that affect spike-ins differently from endogenous RNAs, but further work must clarify this. What is certain is that future work with spike-ins absolutely must keep in mind the compositional nature of the data being generated, and protocols for bulk RNA-Seq may need to be revised to improve the chance of spike-ins accurately anchoring the data to copy numbers.

### Conclusions

In summary, simulating RNA-Seq data using a compositional approach more closely aligns with the kind of data being generated in RNA-Seq. Compositional normalization using negative control features yields a significant improvement over previous methods, in that it performs best in experimental contexts where the composition changes substantially. Importantly, this method can still be safely used in contexts where the compositional changes are unknown. There is much potential to extend the principles of compositional data analysis to other “omics” approaches, since they all generate compositional data; one intriguing possibility is a normalization free method that examines “differential proportionality” [50]. However, our results from simulation and from real datasets demonstrate that, without access to spike-ins or to validated reference features, a researcher is limited in what conclusions can be drawn from *Seq data because of the compositional nature of the data. This work also makes a strong case for there to be more effort to improve and standardize the use of spike-in controls and validated reference features in all “omics” experiments.

## Materials and methods

### absSimSeq approach to simulating RNA-Seq data

See **Fig 1** for a summary diagram of our protocol for **absSimSeq**. When generating RNA-Seq data, the key experimental step which requires a compositional approach is when the actual changes in the RNA content are sampled using an equal but arbitrary amount of RNA by the library preparation process, resulting in a dataset of proportions. To simulate this shift from count data to compositional data, the **absSimSeq** protocol starts with a set of transcripts and their TPMs, either defined by the user or estimated from real data. It then conceptually shifts from considering transcripts per million (a proportion) to considering copy numbers per cell, i.e. the number of transcripts present in each cell (the absolute count unit of interest). It then simulates the fold changes expected to occur between groups directly on the copy numbers, which may or may not result in a substantial change in the total RNA per cell. The next key step is then converting these new expected copy numbers back to TPMs to represent the expected proportion of each transcript that would be present in an equal aliquot taken from each group. These new TPMs are then converted to expected counts per transcript based on their lengths and the desired library sequencing depth, and those expected counts, along with user-defined or estimated parameters for variance within each group, are then submitted to the R package **polyester** to simulate an RNA-Seq experiment.

**AbsSimSeq** also has the option to add spike-ins to the simulated experiment. In our studies, the ERCC ExFold Spike-in mixes are used to define which sequences are included and in what proportions. The user can define what percentage of the transcripts should be coming from spike-ins and which mix to use.

### Simulation of copy numbers for this study

To model a simulated dataset after real data, we took an approach modified from Patro et al [10] and Pimentel et al [13]. To estimate the mean and variance for the control group for our simulation, we wished to use a population without expected biological changes within the group. We thus used as a proxy the largest homogeneous population in the GEUVADIS data set, a set of 58 lymphoblastoid cell lines taken from Finnish women. We estimated transcript abundances using kallisto and human Gencode v. 25 transcripts (Ensembl v. 87), and then estimated negative binomial parameters (the Cox-Reid dispersion parameter) using DESeq2. We next took the mean TPMs from this dataset, for input into **absSimSeq**.

Three simulation studies were performed, with five simulation experiments in each study. The “small” study was intended to simulate experiments where there was no substantial change in the copy numbers per cell per group. The “down” and “up” studies were designed to simulate experiments where there was a large compositional shift, with the total copy numbers either decreasing or increasing.

To simulate differential expression, we first applied a filter where the transcript had to have a TPM value of at least 1. We then randomly and independently assigned each filtered transcript as either not changing (i.e. fold-change of 1) or differentially expressed, using a Bernoulli trial with varying probability of success (5% of all transcripts for the “small” study; 20% for the “down” and “up” studies). For each differentially expressed transcript, a truncated normal distribution was used to simulate the fold change, with a mean of 2-fold, a standard deviation of 2, and a floor of 1.5-fold. A Bernoulli trial was then used to choose either up-regulation or down-regulation with varying probability of success (70% down for the “small” study, chosen to produce roughly equal total RNA in each group; 90% down for the “down” study; 90% up for the “up” study).

The estimated null distribution and the simulated fold changes thus defined the mean copy numbers per cell for the control group and the experimental group, respectively. These copy numbers were then converted back to TPM. Because TPM is proportional to the estimated counts divided by effective length [13], the TPMs were multiplied by the effective lengths and then normalized by the sum to get the expected proportion of reads per transcript per condition. This was then multiplied by a library size of 30 million reads to get the expected reads per transcript per condition. This matrix of expected reads and the Cox-Reid dispersion parameters estimated from the GEUVADIS dataset were used as input for the **polyester** package [32] to simulate 5 samples in each group, with a random variation of about 2-3% introduced into the exact sequencing depth used. The dispersion parameters for the spike-ins was set to the median dispersion of all transcript that had a mean TPM within 5% of the TPM for the spike-in.

**S1 Table** summarizes the simulation parameters and the number of transcripts that are differentially expressed, and **S2 Table** shows the average global copy numbers per cell per condition for each of the fifteen runs. Note that the experimental group in the “up” study had a ∼2.8-fold increase, on average, in the total RNA copy numbers per cell. This is less than the 5.5-fold increase in total mRNA observed after over-expressing the oncogene c-Myc (See the normalized data in Table S2 in Lovén et al., 2012)). The experimental group in the “down” study had a ∼33% decrease in the total RNA copy numbers per cell. This is less than the decrease in total RNA observed in the yeast dataset (See Supplementary Table S2 from [26]).

### Implementing a compositional approach for differential analysis tools: the Log-ratio transformation

To allow tools to use negative control features, like spike-ins or validated reference genes, in a compositional manner, we present a method that uses what is called the “additive log-ratio” (ALR) transformation. This was proposed by John Aitchison to address problems analyzing compositional data. He demonstrated that any meaningful function of compositions must use ratios [29]. Further, he proposed the use of log-ratios to avoid the statistical difficulty arising from using raw ratios. ALR is the simplest of the transformations proposed by Aitchison and others in the field of Compositional Data Analysis. In ALR, if there are D components in a composition, then the D-th component is used as the denominator for all of the other D-1 components.

Formally, if T is a set of D transcripts, then x = {x_t_}_t ⊂ T_ defines the relative abundance of the t-th transcript in the composition, with 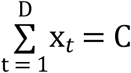, where C is some arbitrary constant (e.g. 1 million for TPMs). These relative abundances are proportional to the units commonly used in RNA-Seq (RPKM, TPM, etc) [18]. The ALR transformation takes a component to be used as the denominator, analogous to the “reference gene” used in qPCR experiments (see “How to interpret the results” below). This forms a new set of D – 1 transformed log-ratios,

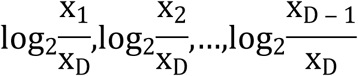

that can then be used for downstream statistical analyses. If one wishes to use a collection of features (e.g. a panel of validated reference genes; a pool of spike-ins), then the geometric mean, g(x), of those multiple features can be used on all D components:

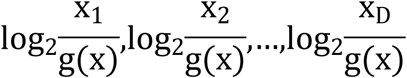

If this is a subset of features, this is called the “multi-additive log-ratio” (malr) [31]. If the geometric mean of all of the features are used, this is the “centered log-ratio” (CLR) transformation introduced by Aitchison and is used in the default mode of ALDEx2 [3]. These are all options available for use in sleuth-ALR.

Log-ratio transformations are undefined if either the numerator or denominator is zero. **Sleuth-ALR** has implemented an imputation procedure for handling zeros that minimizes any distortions on the dependencies within the composition. See Note 5 of **Supporting Information** for more details.

### How to choose a denominator for compositional normalization and how to interpret the results

The proposed interpretation of the results generated by **sleuth-ALR** is simple: whatever the denominator is, the results show how all the other features change relative to that feature or features. For example, if GAPDH is selected as the reference feature, the results show how every other gene is changing relative to GAPDH.

Thus, if there are one or more features which are known *a priori* to be negative controls—a validated reference gene, a pool of spike-ins—these are natural choices for use as a denominator in **sleuth-ALR**, either using a single feature, or using the geometric mean of multiple features. Since the copy numbers of these features are expected to be constant between samples, there is now an anchor for the relative proportions between samples. If negative controls are unavailable, we propose identifying one or more features that have the most consistent proportion across all samples. For **sleuth-ALR**, we chose the coefficient of variation as the metric to measure consistency in proportion. Without external information, though, it is unknown if this “consistent feature” is indeed also consistent in copy numbers. If there is a global change in RNA content, this feature would represent the average change. All current normalization methods assume there is no such change, and so make corrections to the data to remove any perceived change; they are therefore mathematically equivalent to this proposed approach (see [5] for a full mathematical proof). However, this approach has the advantage of making explicit the implicit and necessary interpretation (how are features changing relative to the selected reference feature(s)?). It also provides a feature or set of features that can be used as a reference gene for follow-up validation.

### How sleuth-ALR fits into the current sleuth pipeline

See **S1 Fig** for the pipeline and how it compares to the current pipeline. In the current sleuth pipeline, estimated counts of transcripts from kallisto or salmon are first normalized by the DESeq2 median-of-ratios method [16], and then transformed on the natural log scale with a 0.5-fragment offset to prevent taking the logarithm of zero and to reduce the variability of low-abundance transcripts [13,17]. With the additive log-ratio transformation, the size factor is replaced by the estimated expression of the chosen denominator, and the offset is replaced by the imputation procedure. Once a denominator is chosen, zero values are imputed, the ratios between each feature and the denominator is calculated, and the data is then transformed on a log scale, which can then be used directly in the sleuth model. Our implementation simply replaces the current normalization and transformation functions with the function provided by the **sleuth-ALR** package. For ease of interpretation, the modeling can be done on TPMs directly (using the *which_var* argument for *sleuth_fit*), though the previous choice of modeling the estimated counts can also be used.

### Compositional approach for the other tools

**ALDEx2** has an explicitly compositional approach, and it solves the imputation problem by simulating bootstraps on the data using the Dirichlet-multinomial distribution, which will never yield zero values. It then calculates estimated statistics by examining the differences between groups within each bootstrap. Its default option is to use the CLR transformation, but it has several options for other choices of denominator. A recent paper examined **ALDEx2**’s performance using these different options [31]; and its results suggested that the IQLR (“interquartile range log-ratio transformation”) provided the best balance of performance across real and simulated datasets, with respect to accuracy and computer time. This transformation uses the subset of all features that, after using the CLR transformation, have a sample-wise variance in the interquartile range. Theoretically, this transformation is robust to many features changing either up or down. This and the CLR transformation were used as the normalization methods tested in our study. It can also take a predefined subset of features and uses the geometric mean of those features within each sample; this was the approach taken when utilizing spike-ins for compositional normalization.

**DESeq2** uses the median-of-ratios method [16]. If one wishes to calculate a single size factor to normalize each sample, it is calculated by the *estimateSizeFactors* function. This function has a *controlGenes* option, which allows the user to define a set of features that are expected to have constant expression across all samples. A recent review demonstrated that the DESeq2 size factor is mathematically equivalent to the compositional log-ratio proposed by Aitchison [5]. If a dataset has known negative control features (e.g. spike-ins), these can be used to calculate a DESeq2 size factor similar to what is calculated by **sleuth-ALR** or **ALDEx2**. For **DESeq2**, **edgeR**, and **limma**, we calculated DESeq2 size factors using the *estimateSizeFactors* function with designated negative control features (spike-ins for the simulated data; a validated reference gene for the yeast starvation dataset).

### Pipeline to analyze simulations

The simulated data (FASTA files) were analyzed by **kallisto** for downstream use by all of the tools tested. Spikeins from ERCC Spike-in Mix 1 were included for the simulations (2% of the total RNA), and so were used as the set of features known to have constant expression between samples. Previous studies observed that only highly expressed spike-ins had consistent ratios across samples [33,34]. Thus, we selected spike-ins that had an average log2 concentration of at least 3 between both mixes. This filter results in a set of 47 spike-ins that were used for compositional normalization and for **RUVg** from **RUVSeq**. **RUVg** was used with **DESeq2** and **edgeR** to test its ability to use spike-in information using its own approach.

Filtering is an important issue for managing the accuracy of estimation. Different pipelines make different decisions about what features to filter. To allow the tools to be compared fairly, the same set of filtered transcripts were tested in all tools, defined by those transcripts that passed the standard sleuth filter of having at least 5 estimated counts in at least half of the samples. DESeq2’s default functionality to use independent filtering and Cooks’ outlier filtering did not significantly impact its performance on the simulated data (data not shown), so these were left on.

### Experiments from the original sleuth paper

To see if compositional normalization would produce similar results with fewer replicates, we repeated the self-consistency experiment as described in the sleuth paper [13]. Briefly, we used the Bottomly et al dataset [35], randomly split the 21 samples into a small training dataset consisting of 3 samples in each condition, and a large validation dataset consisting of remaining samples. The “truth” set of features was defined by the hits identified in the larger validation dataset. This was repeated 20 times. At each of three FDR levels (0.01, 0.05, 0.1), we compared the smaller dataset against the larger dataset, and plotted the estimated FDR and sensitivity relative to sleuth-ALR. Since spike-ins were not used in this experiment, and it is unknown if there was any significant change in the total RNA between the groups, the denominator for compositional normalization was chosen based on which feature had the lowest coefficient of variation across the whole dataset. Zfp106-201 (ENSMUST00000055241.12 in Ensembl v. 87) was used as the denominator for **sleuth-ALR** in all datasets. This was also used for RUVg. The IQLR transformation was used for ALDEx2, and otherwise the current normalization methods from the original sleuth paper were used.

To test the performance of compositional normalization when analyzing a negative control dataset, we also repeated the null resampling experiment as described in the sleuth paper [13]. Briefly, we used the Finnish samples from the GEUVADIS dataset, and randomly subsampled the data into twenty null experiments with 3 samples in two groups. This subsampling was stratified by lab to minimize technical variability that may have occurred between labs. Because of the homogeneous population and minimized technical variation, the expectation is that there would be zero differentially expressed features. The null experiments were analyzed, and the number of false positives was plotted at the transcript-level and gene-level. The same denominator was used for compositional normalization in **sleuth-ALR** across all twenty of the null experiments: SRSF4-201 (ENST00000373795.6) for the transcript-level and SRSF4 (ENSG00000116350) for the gene-level. This transcript and gene were determined to have the respective lowest coefficient of variation across all of the samples used for this experiment.

### Pipeline to analyze yeast dataset

To test the different tools and normalization approaches on a real dataset, we chose a well-characterized “yeast starvation” dataset [26]. In this dataset, yeast were cultured in two conditions: (1) freely proliferating using Edinburgh Minimal Medium; (2) the same medium without a nitrogen source (NH_4_Cl), resulting in the cells reversibly arresting into a quiescent state. Two samples from each condition were processed for poly-A selected RNA-Seq or for total RNA (no selection or depletion step). A collection of 49 mRNAs were selected for absolute quantification using the Nanostring nCounter, which uses a fluorescent tagging protocol to digitally count mRNA molecules without the need for RNA purification. The results were normalized to external RNA controls to estimate copy numbers per cell of each mRNA. We used the absolute counts summarized in Supplementary Table S2 of [26] as a basis for selecting opt3 (Pombase: SPCC1840.12.1) as the gene with the smallest coefficient of variation for estimated absolute counts among all samples. This can be considered a validated reference gene. Thus, it was used with methods utilizing negative control features (**sleuth-ALR**, **RUVg**, and other tools using **DESeq2**’s estimateSizeFactors with controlGenes argument) to normalize the data. **Sleuth-ALR** was also tested using rqc1 (Pombase: SPAC1142.01.1), which was selected as having the smallest coefficient of variation for raw abundances (TPM values) across all the samples. This gene represents the “average global trend” or “average global change” in the data, as discussed in “How to choose a denominator” section above.

To re-analyze the RNA-Seq data, we downloaded the *Schizosaccharomyces pombe* genome cDNA FASTA file from ftp.ensemblgenomes.org (Fungi release 37). This was used as the reference for generating the kallisto index. Each tool was then run using default settings.

## Declarations

### Availability of data and code

The yeast starvation dataset was taken from Marguerat et al [26] from ArrayExpress at accession E-MTAB-1154, and the absolute counts were taken from Supplementary Table S2 from [26]. The GEUVADIS Finnish data can be found at ArrayExpress using accession E-GEUV-1, using the samples with the population code “FIN” and sex “female”. The Bottomly et al data [35] can be found on the Sequence Read Archive (SRA) using the accession SRP004777. Human annotations were taken from Gencode v. 25 and Ensembl v. 87, mouse annotations were taken from Gencode v. M12 and Ensembl v. 87, and yeast annotations were taken from Ensembl Genomes Fungi release 37. The code and vignette for **absSimSeq** can be found on GitHub at www.github.com/warrenmcg/absSimSeq, the code and vignette for using sleuth-ALR can be found at www.github.com/warrenmcg/sleuth-ALR, and the full code to reproduce the analyses in this paper can be found at www.github.com/warrenmcg/sleuthALR_paper_analysis. Here are the versions of each of the software used: **kallisto** v. 0.44.0, **limma** v. 3.34.9, **edgeR** v. 3.20.9, **RUVSeq** 1.12.0, and **DESeq2** 1.18.1; the version of **polyester** used is a forked branch that modified version 1.14.1 with significant speed improvements (found here: www.github.com/warrenmcg/polyester); the version of **sleuth** used is a forked branch that modified version 0.29.0 with speed improvements and modifications to allow for **sleuth-ALR** (found here: www.github.com/warrenmcg/sleuth/tree/speedy_fit); the version of **ALDEx2** used is a forked branch that modified version 1.10.0 to make some speed improvements and to fix a bug that prevented getting effects if the ALR transformation with one feature was used (found here: www.github.com/warrenmcg/ALDEx2). All R code was run using R version 3.4.4, and the full pipeline was run using snakemake.

#### Funding

WAM and JYW are supported by the NIH (F30 NS090893 to WAM; R01CA175360 and RO1NS107396 to JYW). HP is supported by the Howard Hughes Medical Institute Hanna Gray Fellowship.

#### Author’s Contributions

WAM conceived the idea, designed the approach, and wrote the software for sleuth-ALR and absSimSeq. WAM and HP wrote the code for the analysis pipeline. JYW and LP provided supervision. WAM and JYW wrote the manuscript.

##### Acknowledgments

We are grateful to Rosemary Braun and David Kuo for helpful suggestions and critical reading of the manuscript.

## Supplemental Information Captions

### Supplemental Notes

#### Supporting Information

Contains Supplementary Notes 1-5. Note 1 argues that *Seq datasets are compositional datasets. Note 2 discusses the three requirements of techniques for analyzing compositional data. Note 3 discusses RUVg and the Compositional Behavior of Spike-ins. Note 4 discusses extending the compositional approach to other high-throughput methods. Note 5 discusses how sleuth-ALR handles zeros.

**Supplemental Figure Legends**

**S1 Fig. The sleuth-ALR approach for compositional normalization.** Under the sleuth model, an observation is modeled as having some error associated with it that is due to the inferential procedure. The true value is modeled as a linear combination of covariates and biological noise. In the original sleuth model (shown in the bottom left), the estimate for the noisy observation was the estimated counts for feature *i* in sample *j*, normalized by the DESeq2 size factor. This and other current normalization methods attempt to translate purely relative information to inferences about absolute changes, but only by assuming no change to the total RNA content. The proposed sleuth-ALR estimate (shown on the bottom right) is an example of how to use compositional normalization. It first focuses on abundances (TPMs) rather than estimated counts, and second normalizes the abundances by a “reference feature”. This avoids having to assume only a few features change, but at the cost of not translating to inferences about absolute changes unless the chosen feature is a validated reference gene or spike-in.

**S2 Fig. A full-range view of the simulation results, accompanying Fig 2.** This shows the full range of FDR and sensitivity for the three simulation studies: **(A)** “small” (5% DE; roughly equal copy numbers in each group); **(B)** “down” (20% DE; ∼33% decrease in copy numbers in the experimental group); and **(C)** “up” (20% DE; ∼2.8-fold increase in copy numbers in the experimental group). This shows that (1) compositional normalization has similar or superior performance throughout the full range of sensitivities and FDR, and (2) RUVg has poor performance, especially when combined with edgeR.

**S3 Fig. ALDEx2 performs similarly in simulations regardless of which statistical method is used**. With the same simulation studies described in **Fig 2**, the performance of ALDEx2 was compared using the Welch t-test (the recommended statistic by the developers), the non-parametric Wilcoxon test, or the reported “overlap” statistic. The overlap statistic is the posterior probability of the effect size being 0 or the opposite direction as what is reported, given the Dirichlet bootstrap samples observed. These three statistics perform approximately similarly no matter which transformation is used: CLR, IQLR, or “denom” (aka ALR, the same as used in sleuth-ALR). This remains true across all three studies: **(A)** “small” (5% DE; roughly equal copy numbers in each group); **(B)** “down” (20% DE; 33% decrease in copy numbers in the experimental group); and **(C)** “up” (20% DE; 2.8-fold increase in copy numbers in the experimental group). This is important because the Welch and Wilcoxon statistics were the ones recommended by the developers but have poor performance when there are few samples.

**S4 Fig. sleuth and sleuth-ALR perform similarly regardless of which statistical method or data unit is used.** With the same simulation studies described in **Fig 2**, the performance of sleuth and sleuth-ALR was compared when using the Wald test or the likelihood ratio test (LRT), as well as when using TPMs or estimated counts for modeling. All combinations perform similarly within each tool across all three studies: **(A)** “small” (5% DE; roughly equal copy numbers in each group); **(B)** “down” (20% DE; 33% decrease in copy numbers in the experimental group); and **(C)** “up” (20% DE; 2.8-fold increase in copy numbers in the experimental group).

**S5 Fig. Spike-ins show a broad range of fold changes and systematic differences in studies with large shifts, accompanying Fig 3.** Using the “ground truth” counts from **polyester,** the log2 fold change was calculated for all spike-ins, and then separated by their concentration in the ERCC Mixes, with “high” expression spike-ins having an average log2 concentration of at least 3 attomoles between both mixes (N = 47 out of 92). These were the spike-ins used for normalization in **Fig 2**. Shown are boxplots of the spike-in fold changes in each experiment across the three studies: (A) “small” (approximately constant total RNA); (B) “down” (∼33 decrease in total RNA); and (C) “up” (∼2.8-fold increase in total RNA). Low-expression spike-ins tend to have a broad range of fold changes, and the high-expression spike-ins tend to have a systematic bias in fold changes in the “down” and “up” studies. For reference, the red dotted line in each run indicates the “ideal” fold change for a spike-in, if it precisely matches the reciprocal of the change in copy numbers between the control and experimental conditions; the blue and gold dotted lines indicate the fold change between conditions of the DESeq2 median-of-ratios and the sleuth-ALR geometric mean of high-expression spike-ins, respectively, suggesting that both are generally good approximations of the “ideal” fold change, and thus are good denominators for normalization.

**S6 Fig. The False Discovery Rate and Relative sensitivity for the Bottomly self-consistency test at additional FDR levels.** This accompanies **Fig 4** in the main text. Shown here are the **(A)** False Discovery Rate, and **(B)** relative sensitivity (% change) at the FDR levels of 0.01 and 0.05.

**S7 Fig. Effect of imputation value on bootstrap variation**. This depicts the summary of bootstraps variation for AAGAB-207 (ENST00000561452.5) within each sample of run #6. The true fold change for AAGAB-207 copy numbers is an 82% decrease. The recommended strategy in Compositional Data Analysis for imputing zero values is to choose a value smaller than the smallest observed value; however, because of the extremely small estimated abundances, this results in a very large variation in the bootstraps within each sample **(A)**. This occurs when at least one bootstrap reports an estimated abundance of zero. Our recommendation is to follow the strategy of previous tools, and choose a larger value to impute. Panel (B) shows the reduction in bootstrap variation after choosing 0.01 for the imputation. The wide variation observed in (A) resulted in a non-significant q-value (0.450), whereas the stabilized variation observed in (B) resulted in a significant q-value (0.047).

**S8 Fig. Effect of imputation on overall simulation performance**. This depicts the full sensitivity versus false discovery rate curve for different choices of imputation value, as compared to standard sleuth as well as the recommended strategy of choosing a value smaller than the smallest observed value (here depicted as “sleuth-ALR counts” for A-C and “sleuth-ALR TPM” for D-F). (**A**) and (**D**) show the results for the “small” simulation group (5% DE; <2% change in copy numbers per cell); (**B**) and (**E**) show the results for the “down” simulation group (20% DE; 33% decrease in overall copy numbers per cell); (**C**) and (**F**) show the results for the “up” simulation group (20% DE; 2.8-fold increase in overall copy numbers per cell). There is improved performance of using imputation versus no imputation, and there are only minor differences in performance in all three studies among any of the choices for imputation values except 0.1 TPM impute value, which is very high (roughly equivalent to a count imputation of 3), in the “up” study.

**Supplemental Tables Legends**

**S1 Table. Summary of Parameters for Simulation Studies.** For each of the fifteen simulation runs, shown are the parameters to establish the number of differentially expressed (DE) transcripts, as well as the number that are up-regulated versus down-regulated. Only transcripts with a TPM of at least 1 were used to simulated differential expression, but the probability of differential expression was determined by the total number of transcripts (∼200K). The proportion of up-regulated transcripts for the “small” study was tuned to result in similar total RNA content in both conditions. The random number generator seed was chosen solely on the basis of yielding consistency in total RNA content across each run within a study. Also shown are the actual number of DE transcripts present in the set of filtered transcripts used by all tools for each run.

**S2 Table. Total RNA Content Per Cell Per Condition for all Simulation Runs.** For each of the fifteen simulation runs, shown are the total RNA copy numbers per cell for each condition. Also reported are the average change in copy numbers between the two conditions for each study.

**S3 Table. Doubling the copy numbers per cell results in the same composition.** Depicted is a toy example of a simple cell with five genes of varying abundances. After an experimental manipulation, each gene has exactly double the copy numbers per cell compared to the control condition. This results in the same relative abundances, and therefore the same composition.

**S4 Table. Spike-in abundances change with large compositional shifts but still accurately capture fold changes.** Depicted is another toy example using the same cell with five genes. In this case, there are large changes in the mRNA genes, but no change in the rRNA gene. Spike-ins added in equal amounts both before and after RNA isolation, show changes in their abundances, and therefore would have changes in the percentage of reads mapping to them. Despite this change in their abundance, spike-ins accurately capture the true fold changes.

**S5 Table. Spike-in abundances change discordantly when non-poly-adenylated RNA changes.** Depicted is another toy example using the same cell with five genes. In this experiment, there is a small change in the rRNA, but no changes to the mRNA genes. Spike-ins were added in equal amounts after RNA isolation, to simulate the protocol used in the zebrafish dataset. Because of the unobserved rRNA change, the spike-ins are affected by the compositional change and show discordant fold changes when compared to the mRNA genes. Normalizing the mRNA genes to the spike-ins results in artefactually elevated fold changes. Thus, the discrepancy observed in the zebrafish dataset can be explained by unobserved changes in the rRNA.

**S6 Table. Spike-ins must be added before RNA isolation to accurately capture true fold changes.** Depicted is a final toy example using the same cell with five genes. In this experiment, there are large and varying changes to both the rRNA and mRNA genes. Spike-ins were added in equal amounts both before and after RNA isolation. Only the spike-in added before RNA isolation can accurately capture the fold changes of the mRNA genes; the spike-in added after is itself affected by the compositional shift of the simultaneous changes in both the rRNA and mRNA genes.

